# Description of the embryonic development in the convict cichlid (*Amatitlania nigrofasciata*)

**DOI:** 10.64898/2026.03.06.710230

**Authors:** Sayaka Matsuo, Rie Kusakabe, Shun Satoh, Kota Kambe, Kazuya Fukuda

## Abstract

We provide a detailed description of embryonic development in the convict cichlid (*Amatitlania nigrofasciata*) from fertilization to hatching at 26 °C, together with a practical staging table anchored to established teleost reference frameworks. Fertilized eggs were obtained by both natural spawning and artificial fertilization. Unfertilized eggs were ovoid and adhesive, surrounded by a chorion and a sticky mucous layer. Early development proceeded, in broad outline, through the teleost sequence of meroblastic discoidal cleavage, blastula, gastrula, segmentation, and organogenesis. The first cleavage occurred at 1.75 hours post-fertilization (hpf), with subsequent cleavages at 30 min intervals, reaching the 64-cell stage at 4.25 hpf. Cleavage up to the 64-cell stage progressed on a timescale broadly comparable to that reported for other cichlids, whereas the interval from the 64-cell stage to early epiboly was relatively short in this species. The high, sphere, and dome stages occurred at 8, 9, and 10 hpf, respectively, with epiboly initiating at the dome stage. At the dome stage, a marginal thickening interpreted as the presumptive embryonic shield became apparent. During early epiboly, the blastoderm showed pronounced spatial heterogeneity: it was consistently thicker and advanced more rapidly on the prospective embryonic axis side, yielding a readily detectable asymmetry. A morphologically distinct embryonic axis became visible at 40–50% epiboly, and epiboly was completed at 28.5 hpf. Notably, somitogenesis began before epiboly completion (first somites at 85–90% epiboly), indicating temporal overlap between late gastrulation and early segmentation. Major organ primordia became apparent during the overlapping segmentation/organogenesis interval, and hatching occurred around 70 hpf. Newly hatched larvae possessed three pairs of adhesive glands. This staging reference enables reproducible developmental sampling and should facilitate future comparative, mechanistic, and experimental work using the convict cichlid.

## Introduction

Recent accumulating evidence indicates that teleost fishes exhibit complex sociality and well-developed social cognitive abilities (Oliveira, 2013; Bshary *et al*., 2014; Salena *et al*., 2021; Bshary and Triki, 2022; Triki *et al*., 2025). Because core neural substrates implicated in these abilities, such as the social decision-making network, are expected to be broadly conserved across vertebrates, including teleost fishes (O’Connell and Hofmann, 2011, 2012; Oliveira, 2013; Bshary *et al*., 2014; Calvo and Schlussel, 2021), teleost fishes, as an early-diverging vertebrate lineage, offer valuable comparative insight into the evolution of social and cognitive abilities.

Cichlids (Cichlidae) have long served as an influential model lineage in evolutionary biology, especially in studies of rapid diversification and adaptive radiation (Kocher, 2004; Henning and Meyer, 2014; Brawand *et al*., 2014; Ronco *et al*., 2021; Svardal *et al*., 2021). Recently, however, cichlids have also gained prominence as experimental systems for social behavior and cognitive research (Maruska and Fernald, 2018; Félix and Oliveira, 2021; Jordan *et al*., 2021; Lein and Jordan, 2021). Many species are readily maintained under laboratory conditions and reliably exhibit a wide range of social behaviors, including pair bonding (e.g., Oldfield and Hofmann, 2011; Cunha-Saraiva *et al*., 2019), parental care (e.g., Cunha-Saraiva *et al*., 2019; Mendes *et al*., 2021), and the establishment of social hierarchies (e.g., Grosenick *et al*., 2007; Gauy *et al*., 2019). These systems also provide tractable paradigms for investigating social cognition and context-dependent decision-making (Culbert *et al*., 2021; Satoh *et al*., 2021; Ferreira *et al*., 2025). Consequently, cichlids provide a promising platform for integrating phenotype-level descriptions of social behavior with proximate mechanistic approaches examining the neuronal, neuroendocrine, developmental, and genetic bases of these traits.

The convict cichlid, *Amatitlania nigrofasciata,* is a Neotropical cichlid with a long history in ethological research. This species forms stable monogamous pairs and provides biparental care, making it a useful model for studies of mate choice (Gagliardi-Seeley *et al*., 2009; Laubu *et al*., 2017), pair formation (Lamprecht and Rebhan, 1997; Gumm and Itzkowitz, 2007; Snekser and Itzkowitz, 2019), and parental roles (Itzkowitz *et al*., 2001; Snekser *et al*., 2014). Recent work has further highlighted its value for addressing emerging questions about prosociality and affective processes (Laubu *et al*., 2019; Satoh *et al*., 2021). Therefore, the convict cichlid has the potential as a teleost model for broadening our understanding of how sociality, cognitive ability, and related traits have evolved across vertebrates.

A key next step is to integrate these behavioral phenotypes with proximate mechanisms. Molecular and genetic toolkits, such as CRISPR/Cas9-based genome editing, have greatly expanded the feasibility of causal tests linking candidate mechanisms to behavioral traits (Alward *et al*., 2020; Sommer-Trembo *et al*., 2024). Applying such approaches to non-traditional models is often facilitated by foundational developmental references. Detailed staging series and developmental timetables are available for widely used model teleosts, such as zebrafish (Kimmel *et al*., 1995) and medaka (Iwamatsu, 2004). In contrast, although morphology-based developmental descriptions are available for several cichlid species (e.g., Meijide and Guerrero, 2000; Kratochwil *et al*., 2015; Mattos *et al*., 2015; Paes *et al*., 2015; Piesiewicz *et al*., 2024), these references are typically species-specific and therefore need to be established for each model species. Despite extensive behavioral research that underscores the convict cichlid’s high potential as an experimental model, a description of embryonic development and a practical staging table have been lacking. Filling this gap would strengthen the utility of the convict cichlid as an integrative model system. Here, we present a continuous description of embryonic development in the convict cichlid from fertilization to hatching under controlled laboratory conditions, together with a morphology-based staging table anchored to widely used teleost reference frameworks. This developmental reference is intended to support future integrative studies using this species.

## Materials and methods

### Maintenance of adult fish

Sexually mature convict cichlids were collected from Okinawa-jima Island, Japan, during 2023–2024 and transported to Kitasato University. Fish were separated by sex and housed in 180 L (90 × 45 × 45 cm) stock tanks. In addition, some fish were maintained in mixed-sex 180 L stock tanks under the same conditions. The water temperature was maintained at 26 °C under a 14 h light/10 h dark photoperiod. Fish were fed a commercial fish food twice daily.

### Collection of fertilized eggs

Although the imminence of spawning could be inferred from several behaviors (Oldfield and Hofmann, 2011) and from female morphological signs (abdominal enlargement and protrusion of the urogenital papilla), the exact timing of spawning was difficult to predict in this species. Thus, we described embryonic development using fertilized eggs obtained by two approaches, as detailed below. As the first, we used naturally spawned eggs. Naturally fertilized eggs were collected as rapidly as possible after spawning and used for observations. Convict cichlids exhibit sexual dichromatism (Noonan, 1983; Beeching *et al*., 1998): sexually mature females show a vivid reddish patch on their abdomen, which is absent in males. In nature, convict cichlids form size-assortative pairs with males tending to be the larger partner (Wisenden, 1995). Based on this knowledge, we selected a female with bright abdominal red and a male approximately 30–50% larger in standard length than the female, and introduced each pair into spawning tanks (40 L; 45 × 30 × 30 cm), from the mixed–sex stock tanks. Spawning tanks were maintained under the same conditions as the stock tanks. As a spawning substrate, one polyvinyl chloride pipe (75 mm diameter and 100 mm length) was provided in each tank. Because females are known to spawn approximately eight days after being introduced into a tank with a potential mate (Santangelo and Itzkowitz, 2004), if spawning did not occur within one week, the pair was returned to the stock tanks and a new pair was introduced into the spawning tank. Typically, females deposited some adhesive eggs onto the substrate progressively, with the male releasing sperm thereafter; the partners alternated gamete release. Immediately after completion of the spawning sequence, the substrate was removed from the tank and the fertilized eggs were collected.

Natural spawning described above yielded a high proportion of well-fertilized eggs but did not allow precise documentation of the earliest cleavage stages. Therefore, as a second approach, we performed artificial fertilization. To avoid using females immediately after spawning, we selected some females from the sex–separated stock tanks, anesthetized them with tricaine methanesulfonate (MS-222; E10521, Sigma-Aldrich) and injected intraperitoneally with human chorionic gonadotropin (hCG; Kyoritsu Seiyaku Co.) at 2 U/g body weight (first dose). Eighteen hours later, the same procedure was repeated with the same dose of hCG plus 17, 20 beta-dihydroxy-4-pregnen-3-one (P6285, Sigma-Aldrich) at 4 μg/g body weight (second dose). Twenty-four hours after the second dose, the abdomens of the females were swollen, but spawning was not observed. In several cichlid species, eggs and sperm have been obtained by gently massaging the abdomens of females and males (Kratochwil *et al*., 2015; Paes *et al*., 2015); however, this method did not yield gametes in the present species. Therefore, in a laboratory kept at 25–26 °C, two females were deeply anesthetized and then euthanized with benzocaine (051-03835, Fujifilm Wako Pure Chemical Co.) 24 h after the second dose, and dissected to remove the ovaries and collect mature oocytes. The unfertilized eggs were transferred into 90 mm petri dishes filled with sterilized hatching buffer at 26 °C (17 mM NaCl, 0.4 mM KCl, 0.27 mM CaClL, 0.65 mM MgSOL, a small amount of methylene blue, and pure water). Also, one male was deeply anesthetized and then euthanized, and dissected to remove the testes. The testes were minced with scissors in a small volume of pure water, and the suspension was examined under a microscope to confirm sperm motility. Several drops of the sperm suspension were then added to the petri dishes containing unfertilized eggs and gently mixed, the eggs were left undisturbed for three min to allow fertilization.

### Observation of embryonic development and image acquisition

For observations of embryonic development, we prepared a working stage. The bottom of a 90 mm petri dish was thinly coated with sterilized 1% agar (010-08725, Fujifilm Wako Pure Chemical Co.), and small holes were formed in the agar using forceps. Fresh hatching buffer was then added to the dish, and fertilized eggs were placed in the holes. While more than 20 fertilized eggs were incubated under identical conditions, six were randomly selected and monitored continuously. During observations, petri dishes were kept in a laboratory maintained at 25–26 °C. Except during photography, dishes were covered with lids. Embryos were observed every 15 min from fertilization to the high stage, every 30 min from the high to dome stages, and thereafter every hour until hatching, and photographed when necessary. Embryonic stages were recorded in hours post-fertilization (hpf). Each developmental stage was defined as the time when ≥50% of embryos from the same batch had reached that stage. For naturally spawned eggs, the exact fertilization time was inferred by aligning stage transitions with those of artificially fertilized eggs. Specifically, we assumed that the time when embryos advanced from the stage at collection (most often the 64-cell stage) to the next stage matched the hpf at which artificially fertilized embryos showed the same transition, and we estimated fertilization time from this value to assign subsequent developmental times. Photographs were taken with a stereomicroscope (SZX12, Olympus) with a digital color camera (Axiocam 208 color, Carl Zeiss). For focus stacking, two to five images taken at different focal planes were combined using Adobe Photoshop (ver. 27.0.0, Adobe Systems).

In this study, the degree of epiboly was quantified as follows: in lateral view, a straight line was drawn between the apices of the animal and vegetal poles of the egg (the longitudinal axis), and % epiboly was defined as the position of the blastoderm margin along this line, expressed as a proportion of the distance from the animal pole to the vegetal pole.

### Ethical Statement

At Kitasato University, there are currently no institutional ethical regulations specifically for research on fish. Therefore, all procedures were conducted in accordance with the Guidelines for the ethical treatment of nonhuman animals in behavioral research and teaching of the Association for the Study of Animal Behaviour and Animal Behavior Society (ASAB Ethical Committee/ABS Animal Care Committee, 2023), and Ethical justification of *Journal of Fish Biology* (2006, 2011).

The convict cichlid was introduced into Japan as an ornamental fish, and individuals that subsequently entered natural waters are thought to have established a feral population on Okinawa-jima Island sometime between approximately 1970 (Tachihara *et al*., 2002) and 1989 (Ishikawa *et al*., 2013). All specimens used in this study were collected from this feral Okinawa-jima population. This study complied with Japanese law and the guidelines of the Ichthyological Society of Japan.

## Results

We described the early development of the convict cichlid from fertilization to hatching. The overall developmental stages were summarized in Table 1. For staging criteria and terminology, we primarily referred to the standard developmental series of zebrafish (*Danio rerio*) (Kimmel *et al*., 1995) and medaka (*Oryzias latipes*) (Iwamatsu, 2004), and also consulted a non-model teleost reference, *Perca fluviatilis* (Alix *et al*., 2015). In addition, we compared our observations with descriptions of Neotropical cichlid development in the Midas cichlid (*Amphilophus xiloaensis*) (Kratochwil *et al*., 2015), *Cichlasoma dimerus* (Meijide and Guerrero, 2000), oscar (*Astronotus ocellatus*) (Paes *et al*., 2015), blue discus (*Symphysodon aequifasciatus*) (Mattos *et al*., 2015), jaguar cichlid (*Parachromis managuensis*), green terror (*Andinoacara rivulatus*), red discus (*Symphysodon discus*) (Piesiewicz *et al*., 2024), and the African cichlid, Nile tilapia (*Oreochromis niloticus*) (Fujimura and Okada, 2007).

**Table 1.**
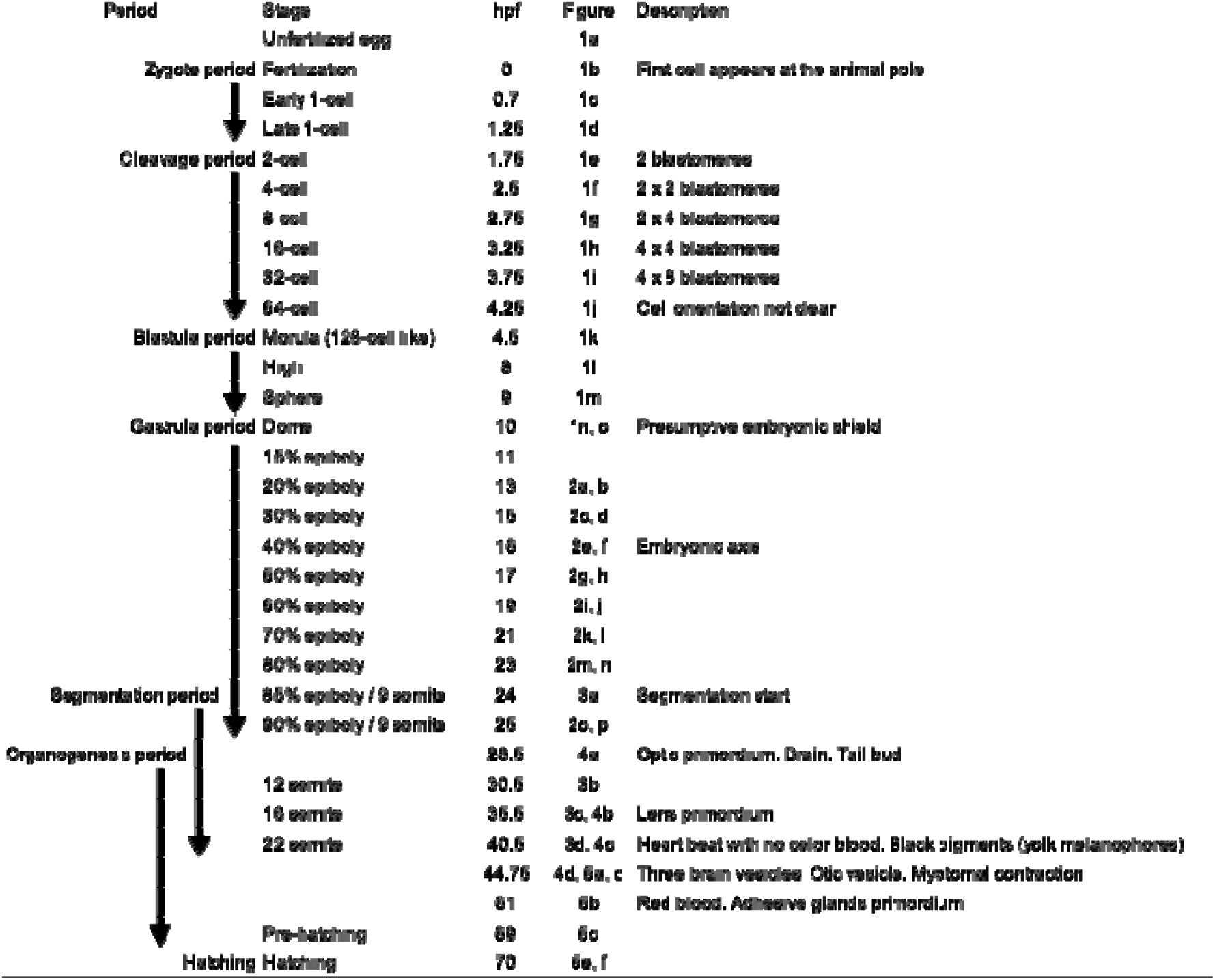
Summary of developmental stages of the convict cichlid, *Amatitlania nigrofasciata*

### Oocyte (unfertilized egg) morphology

Unfertilized eggs (oocytes) were ovoid in shape (Fig. 1a). Each egg was surrounded by a chorion, with a small perivitelline space at the animal pole, where a funnel-shaped micropyle was present. The outer surface of the chorion was further covered by a thin, translucent, sticky mucous layer, which caused the eggs to adhere to the substrate. The yolk mass consisted of semi-transparent, brownish yolk globules, giving the egg a granular appearance.

**Figure 1.**
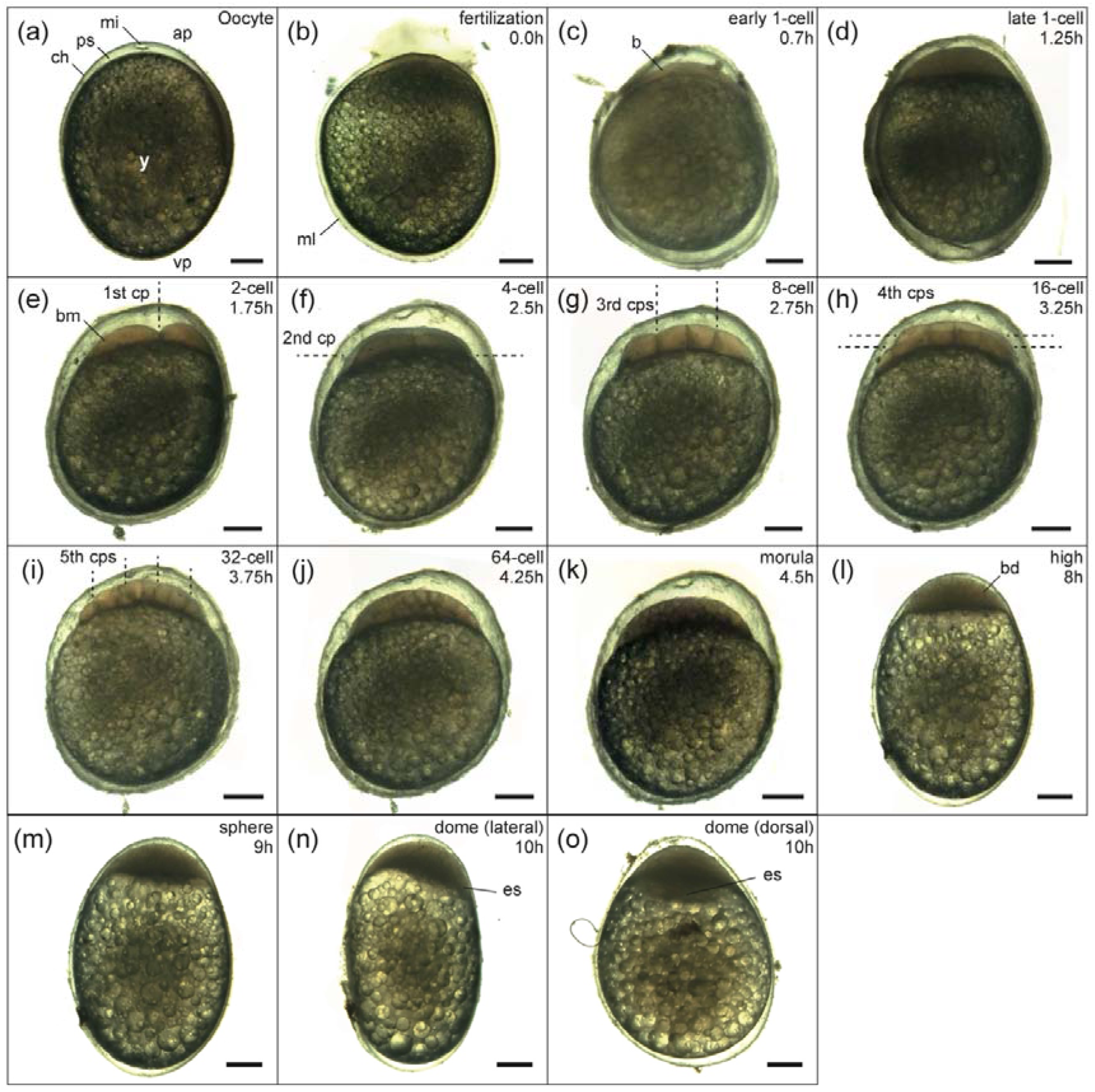
Unfertilized egg and embryos during the zygote, cleavage, and blastula periods. (a) Unfertilized egg. (b) Zygote immediately after fertilization (0 hpf). (c) Early 1-cell stage (0.7 hpf). (d) Late 1-cell stage (1.25 hpf). (e) 2-cell stage (1.75 hpf). (f) 4-cell stage (2.5 hpf). (g) 8-cell stage (2.75 hpf). (h) 16-cell stage (3.25 hpf). (i) 32-cell stage (3.75 hpf). (j) 64-cell stage (4.25 hpf). (k) Morula stage (4.5 hpf). (l) High stage (8 hpf). (m) Sphere stage (9 hpf). (n) Lateral view of the dome stage (10 hpf). (o) Dorsal view of the dome stage (10 hpf). Abbreviations: ap, animal pole; b, blastodisc; bd, blastoderm; bm, blastomeres; ch, chorion; cp(s), cleavage plane(s); es, embryonic shield; mi, micropyle; ml, mucous layer; ps, perivitelline space; vp, vegetal pole; y, yolk. Scale bar = 250 μm

### Zygote period (0–1.75 hpf)

*One-cell stage (0 h)*. Unlike in zebrafish, where the chorion swelled and lifted away from the egg surface after fertilization (Kimmel *et al*., 1995), but similar to the condition described for the Midas cichlid (Kratochwil *et al*., 2015), no clear swelling or separation of the chorion from the egg surface was observed. Fertilization activated cytoplasmic movements, and cortical cytoplasm streamed toward the animal pole (Fig. 1c), where it accumulated to form a dome-shaped blastodisc that gradually segregated from the underlying yolk and became clearly discernible by approximately 1.25 hpf (Fig. 1d). At this stage, the blastodisc cytoplasm appeared uniform and deep brown.

### Cleavage period (1.75–4.5 hpf)

After the first division at 1.75 hpf, subsequent cleavages occurred at intervals of roughly 30 min (range 15–45 min) during the cleavage period. The cleavage mode was meroblastic and discoidal, as in other teleosts. The first six divisions of this period (2- to 64-cell stages) produced stereotyped arrays of blastomeres, as reported previously for teleost embryos (Table 1) (e.g., Kimmel *et al*., 1995; Meijide and Guerrero, 2000; Iwamatsu, 2004). Up to the morula stage, the developmental timetable of the convict cichlid was broadly similar to that reported for cichlids (e.g., Fujimura and Okada, 2007; Kratochwil *et al*., 2015).

*2-cell stage (1.75 hpf)*. The timing of the first cleavage was similar to that reported for other cichlids (e.g., Meijide and Guerrero, 2000; Fujimura and Okada, 2007; Kratochwil *et al*., 2015). At the first division, the cleavage furrow was oriented vertically (meridionally) and split the blastodisc into a pair of equally sized blastomeres that overlaid the yolk surface (Fig. 1e).

*4-cell stage (2.5 hpf).* In the second division, a cleavage furrow running perpendicular to the first cleavage plane split the two blastomeres into four, all of which remained attached to the yolk (Fig. 1f).

*8-cell stage (2.75 hpf).* At the third cleavage, two planes parallel to the first furrow subdivided the four blastomeres, yielding a total of eight cells. In the lateral view, four blastomeres in direct contact with the yolk were clearly visible (Fig. 1g).

*16-cell stage (3.25 hpf)*. During the fourth cleavage, two new furrows formed parallel to the second cleavage furrow, subdividing the two rows of four blastomeres into four rows of four, which remained visible in contact with the yolk (Fig. 1h).

*32-cell stage (3.75 hpf)*. By the 32-cell stage, the cleavage pattern became less stereotyped than in the preceding stages. Because of the increased number of cells and their more irregular geometry, individual blastomeres were more difficult to distinguish than at earlier stages. The fifth set of cleavages occurred along four planes parallel to the first cleavage furrow, subdividing the four rows of four blastomeres into eight rows (Fig. 1i).

*64-cell stage (4.25 hpf)*. After the sixth cleavage, the regular pattern of cleavage planes and cell arrangements was no longer apparent. A superficial layer of blastomeres that did not directly contact the yolk began to form, and the blastodisc started to protrude slightly upward into a dome (Fig. 1j).

### Blastula period (4.5–10 hpf)

The blastula period extended from the morula stage (approximately the 128-cell stage) to the onset of gastrulation. The subsequent high and sphere stages were classified following Kimmel *et al*. (1995). Because a marginal thickening suggestive of the presumptive embryonic shield was already apparent at the dome stage in this species, we considered the dome stage to be part of the gastrula period for staging purposes (see Gastrula period for details). From the morula stage onward, particularly during early epiboly, development in this species proceeded faster than in those species (Table 2).

**Table 2.**
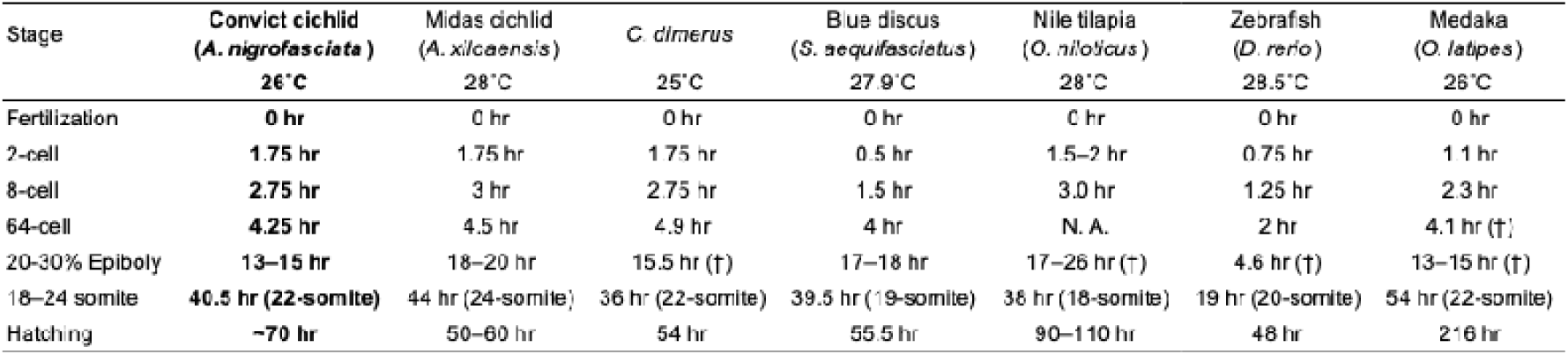
Comparison of developmental rates in some cichlids and established model teleosts (zebrafish and medaka) at the incubation temperatures indicated (convict cichlid: present study; Midas cichlid: Kratochwil *et al*., 2015; *Cichlasoma dimerus*: Meijide and Guerrero, 2000; Blue discus: Mattos *et al*., 2015; Nile tilapia: Fujimura and Okada, 2007; zebrafish: Kimmel *et al*., 1995; medaka: Iwamatsu, 2004). Symbols (†) indicate timings estimated by the authors based on figures and accompanying descriptions in the original publications (i.e., inferred to represent approximately the corresponding stage).

*Morula stage (4.5 hpf)*. Although the exact number of blastomeres could not be counted, the progression of cleavage was evident from the gradual decrease in blastomere size over time. Cleavage planes were no longer regularly oriented, and the blastodisc retained a shallow dome-like shape similar to that at the 64-cell stage, without any apparent increase in its surface area (Fig. 1k).

*High stage (8 hpf)*. Unlike in zebrafish (Kimmel *et al*., 1995), it was difficult in the convict cichlid to determine the number of cells in the blastodisc, nuclei in the yolk syncytial layer, and cells in the enveloping layer. We therefore defined the high stage operationally as the time when the blastodisc was maximally elevated toward the animal pole compared with the preceding and following stages. From this time onward, individual blastomeres were no longer clearly distinguishable, and the cell mass (the blastoderm) appeared as a thickened, multilayered structure (Fig. 1l).

*Sphere stage (9 hpf).* We defined the sphere stage as the time when the area of contact between the blastoderm and the yolk began to expand compared with the preceding stages, and the blastoderm gradually flattened. However, unlike in zebrafish (Kimmel *et al*., 1995), the egg did not become fully spherical, and an ovoid profile remained (Fig. 1m).

### Gastrula period (10–28.5 hpf)

Following the staging criteria for zebrafish (Kimmel *et al*., 1995) and medaka (Iwamatsu, 2004), the onset of the gastrula period was typically demarcated by the emergence of the presumptive embryonic shield region and/or the initiation of involution. However, because we could not resolve the fine-scale cell movements associated with these events (e.g., involution and internalization), we provisionally defined the onset of the gastrula period as the dome stage, when a structure considered to represent a presumptive embryonic shield first became apparent. In contrast to zebrafish, but similar to the Midas cichlid (Kratochwil *et al*., 2015), segmentation began before epiboly was complete in this species.

*Dome stage (10 hpf)*. As described for zebrafish (Kimmel *et al*., 1995), the yolk surface beneath the blastoderm began to bulge upward toward the animal pole, giving it a dome-like appearance (Fig. 1n). At this stage, flattening and spreading of the blastoderm continued, and epiboly had just begun (Fig. 1n, o). From the onset of epiboly, the blastoderm was thicker on the prospective embryonic axis side than on the opposite side (Fig. 1n). In addition, the blastoderm margin on this side advanced toward the vegetal pole slightly earlier than the surrounding margin (Fig. 1o). This marginal thickening, already apparent at this stage, may have corresponded to the earliest manifestation of a presumptive embryonic shield, which functions as the embryonic organizer in teleosts (e.g., Shih and Fraser, 1996).

*Early epiboly stages (20% epiboly, 13 hpf; 30% epiboly, 15 hpf).* At the early epiboly stages, the blastoderm began to spread from the animal pole over the yolk surface. Although epiboly proceeded radially around the animal pole, the rate of progression along the animal–vegetal axis was not uniform; the blastoderm advanced more rapidly on the side corresponding to the embryonic shield (Fig. 2a–d).

**Figure 2.**
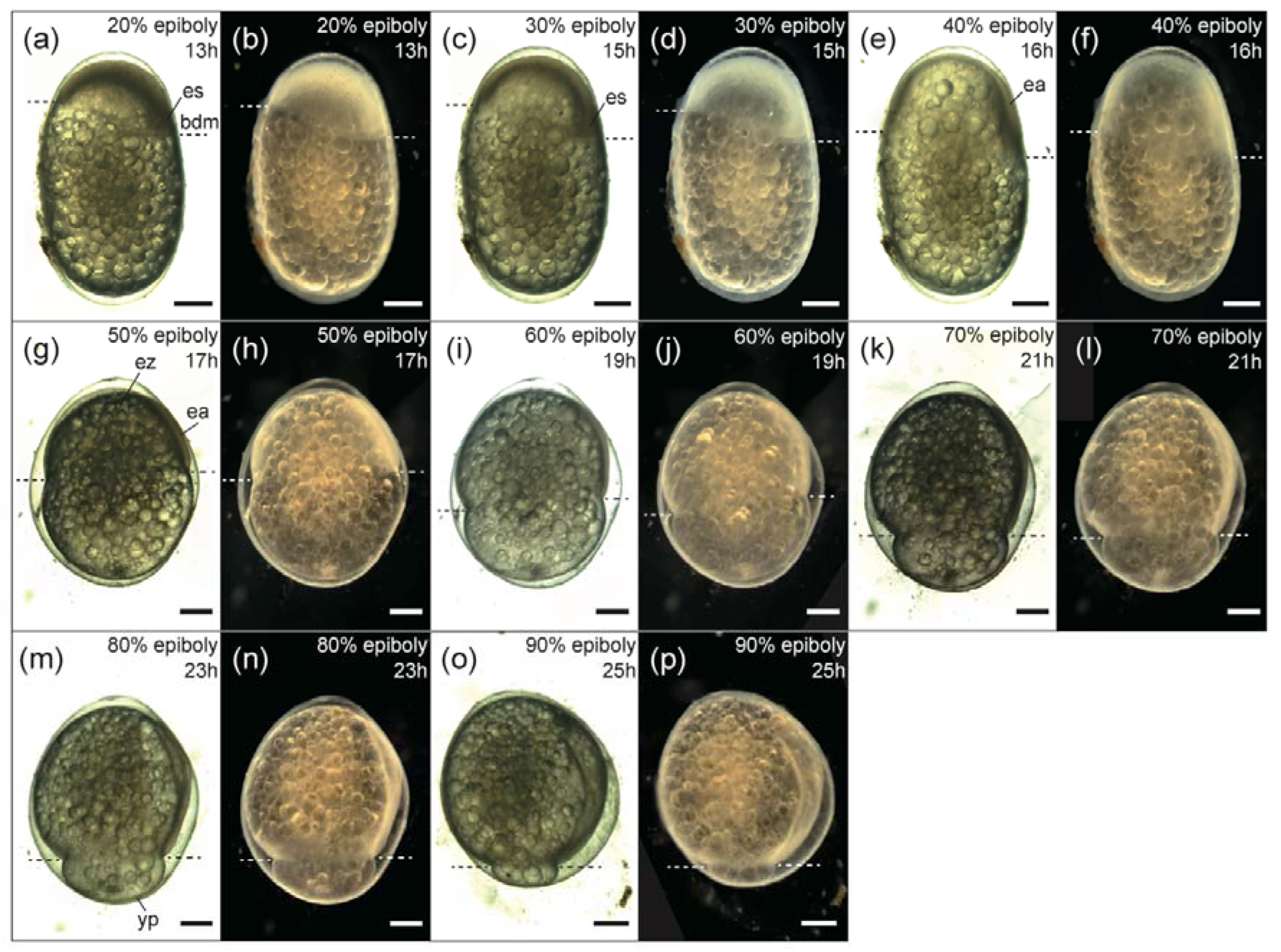
Embryos during the late blastula, gastrula, and early segmentation periods. (a, b) 20% epiboly (13 hpf). (c, d) 30% epiboly (15 hpf). (e, f) 40% epiboly (16 hpf). (g, h) 50% epiboly (17 hpf). (i, j) 60% epiboly (19 hpf). (k, l) 70% epiboly (21 hpf). (m, n) 80% epiboly (23 hpf). (o, p) 90% epiboly (25 hpf). Abbreviations: bd, blastoderm; bdm, blastoderm margin; ea, embryonic axis; es, embryonic shield; ez, evacuation zone; yp, yolk plug. Scale bar = 250 μm

*Middle epiboly stage (40% epiboly, 16 hpf; 50% epiboly, 17 hpf).* Although the progression of the blastoderm margin was radially asymmetric up to 30% epiboly, it gradually became more symmetric around the animal pole as epiboly progressed from 40% to 50% (Fig. 2e–h). During this stage, the thickened cell mass of the embryonic shield became morphologically more distinct and began to elongate along the animal–vegetal axis, beginning to form the embryonic axis (Fig. 2e–h). Epiboly and convergence toward the embryonic axis were accompanied by thinning of the blastoderm over the animal pole, resulting in the formation of an evacuation zone (Fig. 2g, h).

*Late epiboly stages (60%, 19 hpf; 70%, 21 hpf; 80%, 23 hpf; 90%, 25 hpf).* From 50% epiboly onward, the blastoderm margin progressed in a nearly radially symmetrical pattern centered on the animal pole (Fig. 2g–p). In the convict cichlid, somitogenesis began at approximately 85% epiboly (Fig. 3a). Initiation of segmentation before completion of epiboly was similar to that reported for the Midas cichlid (Kratochwil *et al*., 2015) and Nile tilapia (Fujimura and Okada, 2007), but differed from the pattern in zebrafish (Kimmel *et al*., 1995). Epiboly was complete by 28.5 hpf.

**Figure 3.**
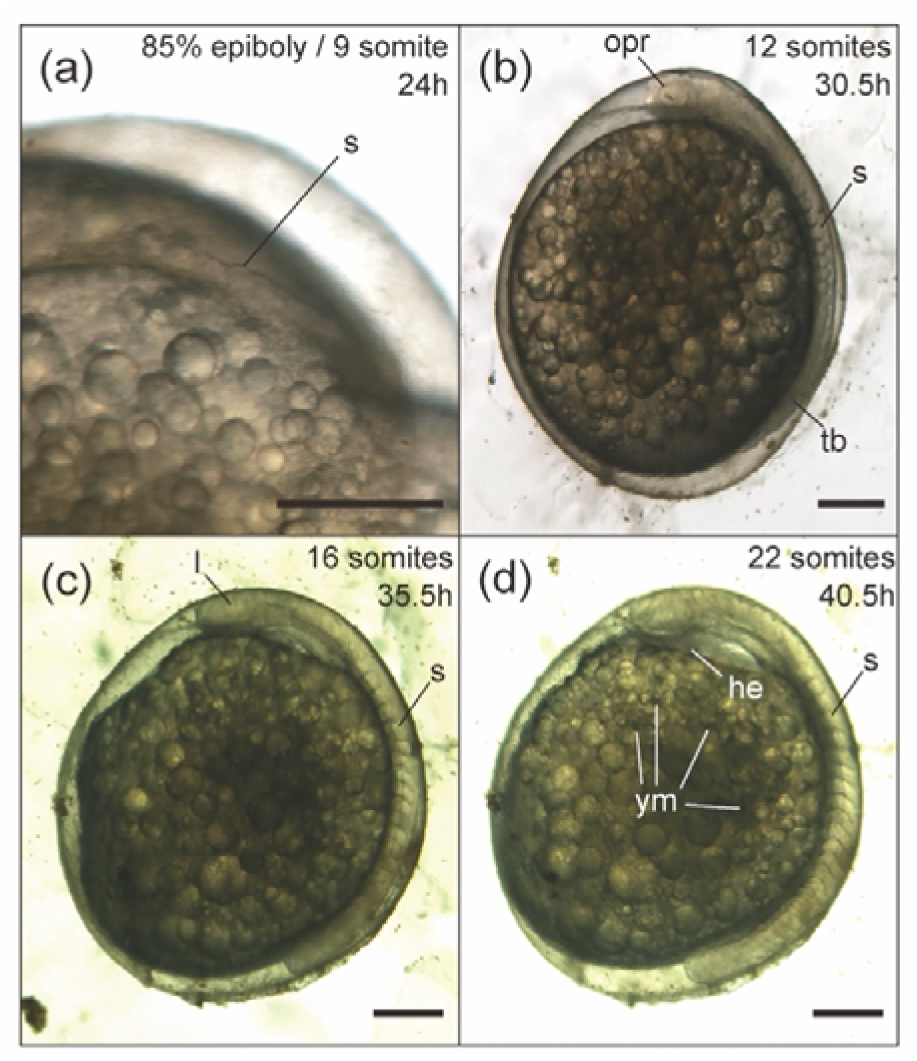
Embryos during the segmentation and organogenesis periods. (a) 85% epiboly (24 hpf), when segmentation first became clearly visible as nine somites under the stereomicroscope. (b) 12-somite stage (30.5 hpf). (c) 16-somite stage (35.5 hpf). (d) 22-somite stage (40.5 hpf). Abbreviations: he, heart; l, lens primordium; opr, optic primordium; s, somites; tb, tailbud; ym, yolk melanophores. Scale bar = 250 μm

### Segmentation period (24–40.5 hpf)

Staging schemes for fish embryos differ among studies and species. In some descriptions (e.g., Kimmel *et al*., 1995; Fujimura and Okada, 2007), the gastrula period is followed by a distinct segmentation period (when somites form) and then a pharyngula period (when pharyngeal arches and many body organs develop), whereas other studies (e.g., Mattos *et al*., 2015; Alix *et al*., 2015; Piesiewicz *et al*., 2024) recognize an organogenesis period after gastrulation, during which primordia of major organs (eye, heart, fins, etc.) become morphologically apparent. In the convict cichlid, somite formation began before epiboly is complete and, as described below, proceeded in parallel with the formation of several major organ primordia. Thus, segmentation and organogenesis could not be separated in time. Nevertheless, to facilitate comparison with other teleost, we recognized both a segmentation period and an organogenesis period that begin during late gastrulation and continue after completion of epiboly, noting that these periods overlap (Table 1). In this study, we operationally defined the segmentation period as extending from the first appearance of clearly distinguishable somites to the stage at which somite number could be reliably counted up to 22 somites under the microscope. During the early part of this period, epiboly was completed, and during the later part, various organogenetic events (such as the onset of heart beating and the appearance of yolk melanophores) were first observed.

*85–90% epiboly / 9-somite stage (24–25 hpf).* At 85–90% epiboly, nine somites were present (Fig. 3a). A tail bud appeared at the posterior end of the body axis. The anterior end displayed the brain primordium and paired optic primordia (Fig. 4a).

**Figure 4.**
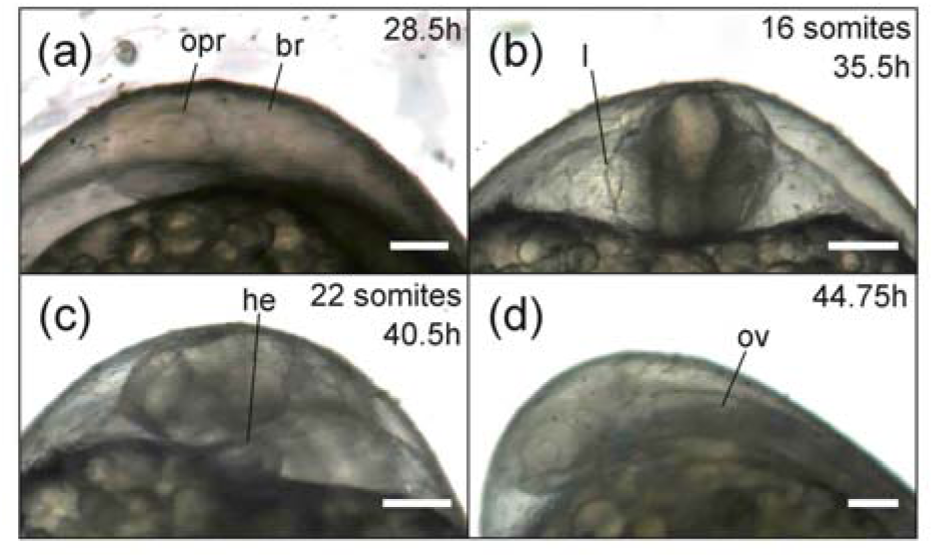
Development of major organ primordia during the organogenesis period. (a) Optic primordium and brain at 28.5 hpf. (b) Lens primordium at 35.5 hpf. (c) Heart primordium at 40.5 hpf. (d) Otic vesicle at 44.75 hpf. Abbreviations: br, brain; he, heart; l, lens primordium; opr, optic primordium; ov, otic vesicle. Scale bar = 100 μm.

*12-somite stage (30.5 hpf).* At the 12-somite stage, epiboly was complete and the blastoderm fully covered the yolk. The optic primordia became prominent. The tail bud extended posteriorly but was not yet clearly prominent (Fig. 3b).

*16-somite stage (35.5 hpf).* At the 16-somite stage, the tail became more distinct and extended. A pericardial cavity formed between the embryo and the yolk, and the lens primordium of the eye became visible (Fig. 3c, 4b).

*22-somite stage (40.5 hpf).* At the 22-somite stage, the lens primordium was easily visible. The first melanophores appeared on the surface of the yolk. The heart primordium first became apparent as a regular contraction of the mesodermal tissue ventral to the head (Fig. 3d, 4c). In most embryos, heartbeats began between 40.5 and 45.75 hpf.

### Organogenesis period (28.5–61 hpf)

As noted above, this period, during which organ formation progresses, overlaps extensively with the segmentation period. In several studies of other fishes, a phase that roughly corresponds to this one has been termed the pharyngula period (e.g., Kimmel *et al*., 1995; Fujimura and Okada, 2007). However, other studies have referred to the same general phase as an organogenesis period (e.g., Alix *et al*., 2015; Mattos *et al*., 2015). In the present study, we referred to this phase as the organogenesis period, to emphasize the broad progression of organ formation during a time window that overlapped with segmentation, rather than the pharyngula period. Morphological changes up to 40.5 hpf were as described in the segmentation period section.

*44.75 hpf.* At 44.75 hpf, myotomal contractions were first observed. The tail continued to elongate along the yolk surface and, after extending around half of the yolk sac, reached the pole opposite the head (Fig. 5a). At this stage, the three primary brain vesicles (forebrain, midbrain, and hindbrain) were morphologically distinct, and a groove marking the midbrain–hindbrain boundary, also known as the isthmic organizer, was visible between the midbrain and hindbrain (Fig. 5a, d). The otic vesicle was forming in the posterior head region (Fig. 4d, 5a). Pigmented melanophores also began to appear on the embryo.

**Figure 5.**
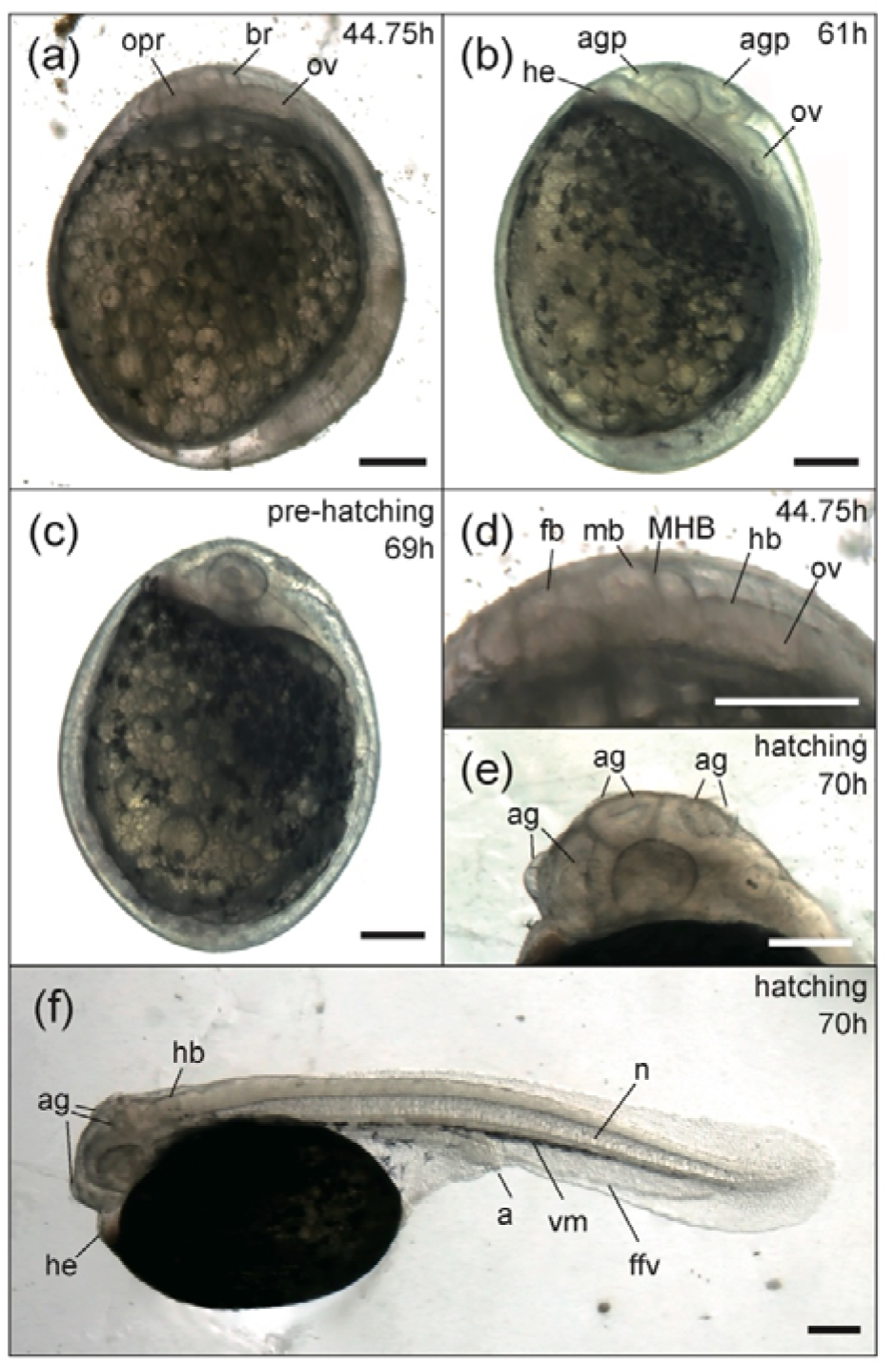
Embryos during the organogenesis period and newly hatched larvae. (a) Embryo at 44.75 hpf. (b) Embryo at 61 hpf. (c) Pre-hatching embryo at 69 hpf. (d) Three primary brain vesicles (forebrain, midbrain, and hindbrain) in a 44.75-hpf embryo. (e) Adhesive glands of a newly hatched larva (70 hpf). (f) Newly hatched larva (70 hpf). Abbreviations: a, anus; ag, adhesive gland; agp, adhesive gland primordium; br, brain; fb, forebrain; ffv, fin fold veins; he, heart; hb, hindbrain; mb, midbrain; MHB, midbrain-hindbrain boundary; n, notochord; opr, optic primordium; ov, otic vesicle; vm, ventral melanophore. Scale bar = 250 μm

*61 hpf.* By 61 hpf, the tail almost encircled the yolk, and embryos frequently showed spontaneous tail flexions within the chorion. The previously colorless circulating blood now appeared red. The brain and otic vesicle were more clearly defined, and melanophores on the embryo became conspicuous (Fig. 5b).

### Hatching (approximately 70 hpf)

Just before hatching (69 hpf), major body organs were clearly visible, and embryos showed frequent body movements (Fig. 5c). The timing of hatching was variable among individuals, occurring within a few hours before or after 70 hpf. Immediately after hatching, the ventral side of the head remained attached to the yolk, and the heart was located anterior to the tip of the snout (Fig. 5f). Three pairs of adhesive glands were present: two pairs were located in the dorsal head region above the midbrain, and one pair was located anterior to the eye (Fig. 5e). After hatching, free-swimming larvae continuously beat their tail from side to side in a regular rhythm.

## Discussion

The present study provided the first continuous description of embryonic development in the convict cichlid from fertilization to hatching, along with a morphology-based staging table. Our observations show that the overall developmental framework conforms to the typical teleost developmental sequence described for other species (e.g., Kimmel *et al*., 1995; Kratochwil *et al*., 2015), progressing from meroblastic cleavage through blastula, gastrula, segmentation, and an organogenesis period (used here, for comparative purposes, as a broad counterpart of the pharyngula period in many teleost developmental tables, such as Kimmel *et al*., 1995), and culminating in hatching at approximately 70 hpf under our rearing conditions. These results demonstrate that convict cichlid embryos share the general developmental organization observed in other teleost species, indicating that this species does not exhibit an unusually derived mode of early development within the broader context of teleost fishes, including cichlids. Nevertheless, detailed species-specific descriptions of developmental timing and morphology remain essential for comparative developmental studies and for integrative approaches, including transcriptomic and other omics-based analyses.

A detailed examination of the developmental process reveals that the convict cichlid shares many characteristics typically observed in other cichlids. The adhesive, ovoid eggs observed in the convict cichlid are consistent with traits reported for several other substrate-brooding cichlids (Meijide and Guerrero, 2000; Paes *et al*., 2011; Kratochwil *et al*., 2015). Notably, we observed the formation of three pairs of adhesive glands (Fig. 5e). These transient larval organs are characteristic of substrate-brooding cichlids and play a crucial role in anchoring the larvae to the substrate before the free-swimming stage (Meijide and Guerrero, 2000; Jones, 1972). The presence and arrangement of these glands in the convict cichlid closely resemble those described in *C. dimerus* (Meijide and Guerrero, 2000) and Midas cichlids (Kratochwil *et al*., 2015), further supporting the conservation of developmental traits associated with the substrate-brooding strategy. In terms of early developmental rates, cleavage progression in the convict cichlid up to the 64-cell stage (4.25 hpf at 26 °C) is broadly similar to that reported for some Neotropical cichlids, although direct comparisons should be interpreted with caution given variation in incubation temperatures across studies (Table 2). In addition, the sequence of organogenesis in the convict cichlid appears largely conserved among cichlids. Key morphological events (e.g., formation of the optic primordium, brain vesicles, otic vesicle, and heart) occur in a similar temporal order to that reported for *C. dimerus* (Meijide and Guerrero, 2000), Nile tilapia (Fujimura and Okada, 2007), and Midas cichlids (Kratochwil *et al*., 2015). These results suggest that the overall schedule of major developmental events in the convict cichlid falls within the range reported for Neotropical and African cichlids, indicating that basic features of the cichlid developmental plan are conserved despite minor heterochronic shifts.

Within this broadly conserved developmental plan, our staging series identifies several features of early development that are characteristic of the convict cichlid compared with established model teleosts, such as zebrafish and medaka, and with other described cichlids. Firstly, the convict cichlid exhibits a relatively short interval between late cleavage and the onset of epiboly compared to other cichlids (Table 2). In the convict cichlid, early cleavage up to the 64-cell stage proceeds at a broadly comparable rate to that reported for other cichlids. By contrast, the interval from the 64-cell stage to early epiboly appears relatively short. In the present study, embryos reached 20–30% epiboly within 8.75–10.75 hours after the 64-cell stage, whereas longer intervals have been reported for the Midas cichlid (13.5–15.5 hours) and blue discus (13–14 hours), although incubation temperatures differ among studies (Table 2). In contrast to this comparatively rapid progression, the time to hatching in the convict cichlid (around 70 hpf) occurs later than in several other cichlids (e.g., Midas cichlids, 50–60 hpf; *C. dimerus*, 54 hpf; blue discus, 55.5 hpf) (Table 2). Overall, these observations suggest that minor heterochronic shifts occur even among closely related cichlids, with species differing in how developmental time is allocated across early versus late phases. A second feature of the convict cichlid is the spatial heterogeneity of blastoderm morphology and movement during early epiboly. In this species, from the onset of epiboly (i.e., the dome stage), the blastoderm was consistently thicker on the side that subsequently gave rise to the embryonic axis, and the blastoderm margin advanced more rapidly on this side during early epiboly. This produces a readily detectable asymmetry. Similar anisotropies in blastoderm organisation or epiboly dynamics have been reported in other cichlids, including localized thinning of the blastoderm during epiboly in *C. dimerus* (Meijide and Guerrero, 2000) and differences in epiboly progression among egg regions in Nile tilapia (Fujimura and Okada, 2007). However, in the convict cichlid, this asymmetry provides an obvious external cue to axis orientation that is visible under a stereomicroscope well, potentially facilitating consistent staging and experimental sampling during early gastrulation. In the present study, we provisionally considered the marginal thickening at the dome stage to represent the presumptive embryonic shield. Because this thickening was continuous with subsequent embryonic axis formation, the organizer region corresponding to the embryonic shield may be established relatively early in this species. The embryonic shield formation and associated cell-layer movements (including involution/internalization) are commonly used to demarcate the onset of gastrulation in teleost developmental tables (Kimmel *et al*., 1995; Iwamatsu, 2004). It should be noted, however, that some developmental descriptions in other cichlids place the start of the gastrula period at later stages, typically around 10–50% epiboly (Fujimura and Okada, 2007; Kratochwil *et al*., 2015; Mattos *et al*., 2015). Reconciling these differences will require higher-resolution analyses of cellular dynamics during early development across cichlids, so that the onset of gastrulation can be defined based on cell movements rather than on external morphology alone. Third, the convict cichlid exhibits temporal overlap between epiboly and the initiation of somitogenesis. In our observations, segmentation began while epiboly was still incomplete (the first somites appeared at around 85–90% epiboly), indicating that late gastrulation and early segmentation occured concurrently. This pattern is similar to that observed in other cichlids, such as Midas cichlids (Kratochwil *et al*., 2015) and Nile tilapia (Fujimura and Okada, 2007), but differs from the developmental partition typically presented for zebrafish (Kimmel *et al*., 1995) and medaka (Iwamatsu, 2004), in which segmentation follows the completion of epiboly. These features emphasise that cichlids can exhibit diversity in the timing and spatial dynamics of early development within a conserved teleost framework. This highlights the importance of morphology-based staging schemes, rather than relying only on time-based developmental criteria.

Beyond providing a developmental description, the staging framework presented here enhances the experimental utility of the convict cichlid. Expanding standardized developmental references for the Neotropical cichlid lineage should facilitate interspecific comparisons of developmental timing and morphology within an adaptive-radiation framework. The convict cichlid has also been widely used in behavioral research, including ethology, social decision-making, and cognitive abilities (e.g., Lamprecht and Rebhan, 1997; Itzkowitz *et al*., 2001; Gumm and Itzkowitz, 2007; Gagliardi-Seeley *et al*., 2009; Snekser *et al*., 2014; Laubu *et al*., 2017, 2019; Snekser and Itzkowitz, 2019; Satoh *et al*., 2021). Our developmental reference should facilitate the application of approaches focused on the proximate mechanisms underlying these behaviors, thereby extending existing phenotype-based behavioral research in this species. In particular, standardized staging enables developmentally matched sampling for neuronal and molecular analyses and supports reproducible timing of experimental manipulations. Because the stages defined here rely on external landmarks that can be readily identified under a stereomicroscope and are comparable to those of widely used reference species (zebrafish, medaka, and Nile tilapia), they should allow straightforward alignment of developmental sampling across taxa. Thus, the present developmental reference should support future comparative studies, as well as experimental embryology and molecular genetic approaches, using the convict cichlid.

Finally, further studies are needed to complement the present study and achieve a more comprehensive understanding of early development in this species. The staging scheme presented here is primarily based on bright-field stereomicroscopic observations of whole embryos. While this approach allowed us to define a continuous series of external morphological landmarks from fertilization to hatching, it has limitations in resolving tissue-and cell-level events. For example, detailed cell movements during gastrulation and the morphogenesis of internal structures, such as the neural tube and gut, could not be fully characterized based on external observation alone. To describe these processes in more detail, complementary approaches involving histological sectioning and staining of fixed specimens will be necessary to correlate internal histological changes with the external stages defined here. Furthermore, the present study was conducted under a single temperature condition (26 °C). The significant influence of incubation temperature on developmental rates in teleost is well recognized (e.g., Kratochwil *et al*., 2015). Therefore, to facilitate robust comparisons with related species and to further establish the convict cichlid as a flexible experimental model, future research should examine developmental rates at various temperatures.

## Acknowledgments

We are grateful to the members of the Laboratory of Reproductive Physiology of Aquatic Organisms, Kitasato University, for their assistance during our study.

## Author contributions

Conceptualization: KF; Methodology: SM, RK, SS, KK, KF; Formal analysis and investigation: SM, KF; Writing - original draft preparation: SM, KF; Writing - review and editing: RK, SS, KF; Funding acquisition: SS, KF; Resources: SS, KF; Supervision: RK, KF

